# Oral-stomach sampling to replace rumen-fistulated animals in ruminant nutrition research - a case study

**DOI:** 10.1101/2025.01.17.633359

**Authors:** A. Boudon, D. Kwiatkowski, V. Niderkorn, E. Forano, P. Nozière, M. Silberberg

## Abstract

Studies using fistulated ruminants have shed light on digestion and fermentation in the rumen and yielded tools to determine the feed efficiency of diets, limit effluents or anticipate variability in the quality of animal products. However, the ethical acceptability of such studies has been called into question. Our objective was to determine whether oral-stomach sampling (OSS) is an acceptable alternative to sampling through a ruminal cannula in characterizing the variability of rumen fluid composition induced by an acidogenic dietary challenge in dairy cows. During three four-week periods (P1, P2, P3), six rumen-fistulated cows were fed a standard diet based on supplemented corn silage (periods 1 and 3) alternating with a diet enriched with starch (period 2). Rumen juice was collected through cannulas at three locations in the rumen (reticulum, ventral sac or a mix of both) and by OSS, once per week at 0830 h, i.e. before morning feeding, and once every third week of each period at 1330 h, i.e. 4.5 hours after morning feeding. Whatever the sampling method or location, ruminal pH was lower in period 2 compared to periods 1 and 3 at 1330 h (6.21 vs. 5.57 vs. 6.12 in periods 1, 2 and 3). Ruminal pH was higher when obtained by OSS rather than cannulation, whatever the sampling location (on average, +0.44 points at 0830 h and +0.56 points at 1330 h). Mineral composition indicated a presumed dilution by saliva of OSS samples. This was also consistent with lower concentrations of volatile fatty acids. The next steps will be to analyze the associated variations in in vitro rumen fermentation parameters and microbiota composition.

**Implications:** The use of fistulated ruminants to elucidate ruminal physiology is currently controversial in society. In order to minimise, reduce or replace these experimental models, alternatives to ruminal fistula sampling are being evaluated. Here we focus on oral stomach sampling (OSS). We show that OSS can be a satisfactory alternative to cannula sampling in the evaluation of rumen fermentation parameters, especially for the molar proportion of volatile fatty acids, rumen pH, and rumen volatile fatty acid and ammonia concentrations. We also show that OSS must be standardized to avoid salivary contamination affecting the quality of the results.

## Introduction

For almost a century, rumen-fistulated ruminants have been used in studies that have helped predict the nutritional value of feeds or diets and establish the principles of diet formulation in ruminants (Mould, 2002; Doreau, 2008; Phillips, 2021). These inputs allowed the formulation of diets that optimized animal production performances while maintaining animal health and limiting effluents in the environment (INRA, 2018). One reason for the almost widespread use of this type of animal throughout the world is the ease with which these animals can be kept in a herd once the surgical phase is over (Durand et al., 2021). This is related to the very low risk of infection since the rumen wall is sutured to the skin, and the apparent absence of disturbance to the animal, which can keep it for several years (Doreau, 2008).

However, the use of these animals is controversial (Pagella et al., 2018; Phillips, 2021). This controversy is all the greater when one considers that the aim of the studies in which these animals are involved is not specifically related to animal health or welfare, since an associated aim of these studies is most often to increase animal productivity or efficiency of resource use. Beyond the specific case of rumen-fistulated ruminants, there is also a debate regarding the use of experimental animals, especially in Europe, and a pressing need for replacement, reduction and refinement (MacArthur Clark, 2018; Veissier and Deiss, 2022). However, climate issues are making it increasingly important to increase the feed efficiency of farm animals, reduce enteric methane emissions and characterize the nutritional value of new food resources in the context of agrobiology. It is therefore more important than ever to reach a consensus on the validity of alternatives to rumen-fistulated cows (Mould, 2002; Doreau, 2008; Pagella et al., 2018).

Several types of measurements require rumen-fistulated cows, such as the *in sacco* measurement of ruminal degradability of feeds, measurement of digestive transit, inoculation of in vitro fermenters, dynamic characterization of the physico-chemical properties of rumen contents or microbiota analysis. For the last three types of measurements, direct access to the ruminal content is required. A proven alternative to ruminal sampling through the rumen fistula is oral-stomach sampling (OSS).

Numerous publications have already shown that this technique can lead to measurement bias compared to the reference method of ruminal sampling through the fistula (Geishauser and Gitzel, 1996; Duffield et al., 2004; van Gastelen et al., 2019; de Assis Lage et al., 2020). This was shown to be particularly true for ruminal pH measurements and, to a lesser extent, for ruminal volatile fatty acid (VFA) concentrations and composition. The bias was always a dilution with OSS, either due to contamination by saliva when the probe crosses the oesophagus (Duffield et al., 2004) or to a sampling site in the cranial dorsal rumen rather than in the central ventral rumen because the probe did not cross the fibre mat in the rumen (Shen et al., 2012). However, some authors found that the bias could be limited by optimizing the design of the probe, by adjusting the length of the tube to control the sampling site or by discarding the first 200 mL or more potentially contaminated with saliva (Duffield et al., 2004; Shen et al., 2012). There is also some consensus that OSS can provide a relative measurement, even if a bias is observed.

However, in addition to the need to assess the animal welfare implications, generalization of the use of OSS and abandonment of rumen fistula sampling require proper assessment of the biases inherent in OSS and the consequences this may have for the construction of future experimental designs. None of the above comparisons between OSS and rumen fistula sampling were based on repeated sampling as part of an experimental trial designed to induce specific variability in the measured parameters. Thus, our objective was to validate an optimized OSS method for its ability to highlight differences between treatments in a model experiment to study the effect of the inclusion of an acidogenic diet on the dynamics of ruminal composition in lactating dairy cows. Further work will also compare OSS and cannula sampling in rumen microbiota analysis and in the capacity of rumen juices to inoculate fermenters for test gas.

## Material and methods

The present study took place at IE PL, INRAE, Dairy nutrition and physiology (IE PL, 35650 Le Rheu, France; https://doi.org/10.15454/yk9q-pf68, protocol L2018) between January 4th and March 26th 2020. The experimental design and the experimental procedures were approved by a Regional Ethics Committee and the French Ministry of Research under number APAFiS #26894-2020081715322100_v2.

### Animals and diets

Six rumen-fistulated multiparous Holstein cows in early lactation were involved in this longitudinal study. At the beginning of the experiment, cows were at 93 ± 10 days in milk, they were producing 35.0 ± 12.2 kg milk /day, and weighed 686 ± 72.4 kg. All cows were housed in a barn where each animal was kept in a tie-stall with free access to water and pure salt blocks. Each stall was 1.4 x 2.0 m wide, and was equipped with a rubber mat. Prior to the experiment, all cows were habituated to this environment for one week and maximum ingestion capacity was estimated for each individual by insuring 10% feed refusals over this period on a standard milk production diet.

During the experiment all cows received successively 3 diets distributed as total mixed rations (TMR) (Table 1) each for 4 weeks. During the first experimental period (P1), cows received the same control diet as during the pre-experimental week which was the regular milk production diet of the experimental facility. This diet contained a low amount of concentrate (12% DM) that provided a low amount of starch (18% starch DM). During the second period of the experiment (P2), cows received a challenging diet corresponding to a high-energy diet (55% DM of concentrate) that provided a high starch content (29% DM). This high starch diet was progressively introduced to cows over 10 days to avoid acute rumen acidosis. Although this high-energy diet is classically used to maximize milk production, it represents a risk of SARA (subacute rumen acidosis) due to the high amount of available starch. For the last period (P3), cows could recover with the control diet distributed at the beginning of the experiment after 2 days of progressive transition. Cows received diets twice every day at 0900 h (60% TMR) and at 1700 h (40% TMR). They had free access to diets for 4h each morning and 5h each afternoon to insure a total ingestion of the distributed feed. TMR refusals were weighed every day so that the DMI was calculated individually for the three experimental periods.

**Table 1.** Composition of the diets given to the cows (the diets were fed *ad libitum* as a total mixed diet; the standard diet was fed at periods 1 and 3, i.e. between weeks 1 to 4 and 9 to 12; the acidogenic diet was fed at period 2, i.e. between weeks 5 to 8).

|  | Standard diet | Acidogenic diet |
| --- | --- | --- |
| Feed, g/kg DM |  |  |
| Maize silage | 65 | 31 |
| Soybean meal | 13 | 13 |
| Energy concentrate<sup>1</sup> | 12 | 55 |
| Dehydrated alfalfa | 9 | 0 |
| Mineral feed<sup>2</sup> | 1 | 1 |
| Chemical composition, g/kg DM |  |  |
| Crude protein | 144 | 155 |
| Starch | 216 | 258 |
| NDF | 390 | 296 |
| ADF | 235 | 150 |
| Nutritive value<sup>3</sup> |  |  |
| UFL/kg DM | 0.83 | 0.96 |
| RPB g/kg DM | 0.82 | 2.76 |
| Protein truly digestible in the intestine (PDI, g/kg DM) | 92 | 103 |

Abbreviations: DM = Dry Matter; NDF = Neutral Detergent Fiber; ADF = Acid Detergent Fiber; UFL = "Unité Fourragère Lait", 1 UFL = 7.634 MJ NE<sub>L</sub> (Net Energy for Lactation), RPB = Rumen Protein Balance.
<sup>1</sup> Concentrate consisting of, DM basis, 20.0% wheat, 19.9% corn, 20.0% barley, 15.1% wheat fine bran, 20.4% dried sugar beet pulp, 1.1% rapeseed oil, 2.5 % cane molasses, 1.0% salt.
<sup>2</sup> Mineral feed formulated to supply, DM basis, 5.5% P, 26% Ca, 4% Mg, 0.1 %Na, 1200 mg/kg Cu, 4530 mg/kg Zn, 3500 mg/kg Mn, 80 mg/Kg I, 35 mg/kg Co, 22 mg/kg Se, 500000 IU/kg vitamin A, 100000 vitamin D3, 1500 IU/kg vitamin E.
<sup>3</sup> Nutritive value calculated according to INRA (2018) according to the observed average feeding level of 3 kg DM/100 kg BW considering a total mixed diet fed *ad libitum* to a cow with a body weight of 686 kg, at 120 days in milk with a potential peak milk yield of 40 kg/d, with milk protein content of 32 g/kg and a milk fat content of 40 g/kg.

### Sampling procedures and measurements

#### Feed

Each individual feed as well as the TMR were sampled every week and stored at - 20°C until analysis. Analyses of the nutritional value of feeds were performed by the Dumas method for crude protein on a Leco apparatus (Leco Corp., St. Joseph, MI), by the Van Soest method with a Fibersac extraction unit (Ankom Technology Corp., Fairport, NY) for the fibre contents and by polarimetry according to the European Commission regulation no. 15/2009 for starch content. Those analyses were performed by an external laboratory (Labocea, Ploufragan, France).

#### Milk yield and composition

Animals were milked twice daily (0645 h and 1700 h) and individual milk yield was recorded at each milking. Milk fat and protein contents were determined over 2 successive morning and afternoon milkings each week by infrared spectroscopy by an external laboratory (MyLab, Chateaugiron, France).

#### Reticulo-rumen liquor sampling procedures

Reticulo-rumen liquor was sampled through a cannula using 3 different methodologies: (i) directly in the reticulum to obtain RCn (Reticulum through Cannula samples), (ii) in the ventral sac to obtain VSCn (Ventral Sac through Cannula) samples, and (III) as a mix of reticulum liquor and ventral sac liquor (50/50, v/v) to obtain MCn (Mixed through Cannula) samples. Those samples were obtained by introducing through the cannula a hollow cane connected to a force-feeding gun for calves that provided enough suction to drain juice from the reticulo-rumen or the ventral sac fibrous content. The hose diameter was 20 mm and the inserted length was 200 cm with a stainless steel strainer at the tip. The samples were obtained for each cow once a week at 0815 h, 1330 h and 1630 h. Reticulo-rumen liquor was also obtained via OSS (Popova et al, 2022) via a dedicated drenching/sampling pump (Ammerlaan Import, Longnes, France) shown in Figure1. OSS was performed on each cow once a week at 0845 h on the same day as for cannula sampling. On P1-week 3, P2-week 3 and P3-week 3 a second sampling was performed at 1330 h so that cows were sampled twice a day only 3 times during the whole experiment. For each sampling, in order to avoid salivary contamination, the first 200 mL of rumen liquor was systematically discarded.

The pH of all reticulo-rumen samples was extemporaneously measured using a laboratory pH electrode (Electrode pH - SI Analytics - BlueLine 25 pH) connected to a pH meter (pH3310, WTW, Xylem Analytics, Weilheim, Germany). Samples were then filtered through a nylon fabric (400 µm) and strained rumen fluid was stored at - 20°C in 2 mL Eppendorf tubes after freezing in liquid nitrogen for later mineral concentration analysis. Only samples collected at 0830 h every week and at 1330 h every third week of each period were used for VFA and NH3 analyses. For minerals (Ca, K, Mg, Na, P), only samples collected at 0830 h and 1330 h every third week of each period were analyzed. For VFA analysis, 800 μL of rumen fluid was added to 500 μL of 0.5 N HCl containing 2% (w/v) metaphosphoric acid and 0.4% (w/v) crotonic acid. For NH3 analysis, 500 µL of rumen fluid was added to 50 µL of 5% H3PO4. All sample tubes were stored at -20°C until laboratory analyses. Gas chromatography analyses were performed as already reported by Morgavi et al (2003) for VFAs and spectrophotometry analyses were performed as reported by Morgavi et al (2008) for NH3 and by ICP-OES for Ca, K, Mg, Na, P after dilution of samples with nitric acid (2% v/v).

### Statistical analyses

All data processing and statistical analyses were performed using R3.2.3 software associated with the following packages: ggplot2, lubridate, tidyr, dplyr, lmerTest and emmeans. All the variables collected daily or several times a week throughout the experiment (DM intake, milk production, milk fat and protein contents, rumen pH) were averaged per cow and per week for the plotting of figures. To analyze the influence of the sampling method on the measurements, as well as the dynamics of adaptation of the animals to the change in diet, the statistical individual was the average of the parameters analyzed during the last two weeks of each period for each cow (n=18 cow x period). The statistical model was a mixed model including the fixed effects of the period (P1 vs. P2 vs. P3), the sampling method or location (OSS vs. RCn vs. VSCn and vs. MCn) and the interaction between both, and the random effect was the animal. Differences were considered significant when P < 0.05 and trends were considered when P < 0.10. Post hoc analyses were performed using Tukey’s test to compare adjusted means by period and/or location/method of measurement. Root mean square prediction error (RMSPE) was calculated between OSS (predicted value) and VSCn sampling, and was decomposed into the error of central tendency (ECT), the error due to regression (ER), and the error of dispersion as described by Bibby and Toutenburg (1977). VSCn was considered to be the best method to obtain a mean representative sample of the whole rumen during repeated sampling because it is supposed to be less subject to immediate and transient dilution by saliva during eating and ruminating mastication (Shen et al., 2012).

## Results

Two of the six animals involved in the experiment presented bedsores and hoof inflammation at the very beginning of P2 (high starch period). These clinical signs disappeared rapidly and so we kept all animals in the experiment.

### Ingestion and milk composition

Average DMI, milk production, milk fat and protein contents were 20.1 ± 1.92, 36.1 ± 5.61, 26.8 ± 8.06 and 28.3 ± 2.79 g/kg, respectively, throughout the experiment (average between weeks 1 to 12, Figure 2). Considering the average of the last two weeks of each period, DMI was unaffected by the period whereas milk production was higher in P2 compared to P1 and P3 (39.2 vs. 34.6 kg/d, P < 0.05). Milk fat content decreased between P1 and P2 from 33.1 to 15.8 g/kg and increased in P3 to 29.0 g/kg, but remained lower than in P1 (P < 0.001). Milk protein content was slightly higher in P2 than in P1 or P3 (P < 0.01). Like the milk fat content, the ratio between milk fat and protein contents decreased from 1.18 in P1 to 0.54 and P2 and increased at P3 to a value of 1.04, which was significantly lower than that of P2 (period effect P < 0.001).

**Fig. 1.**
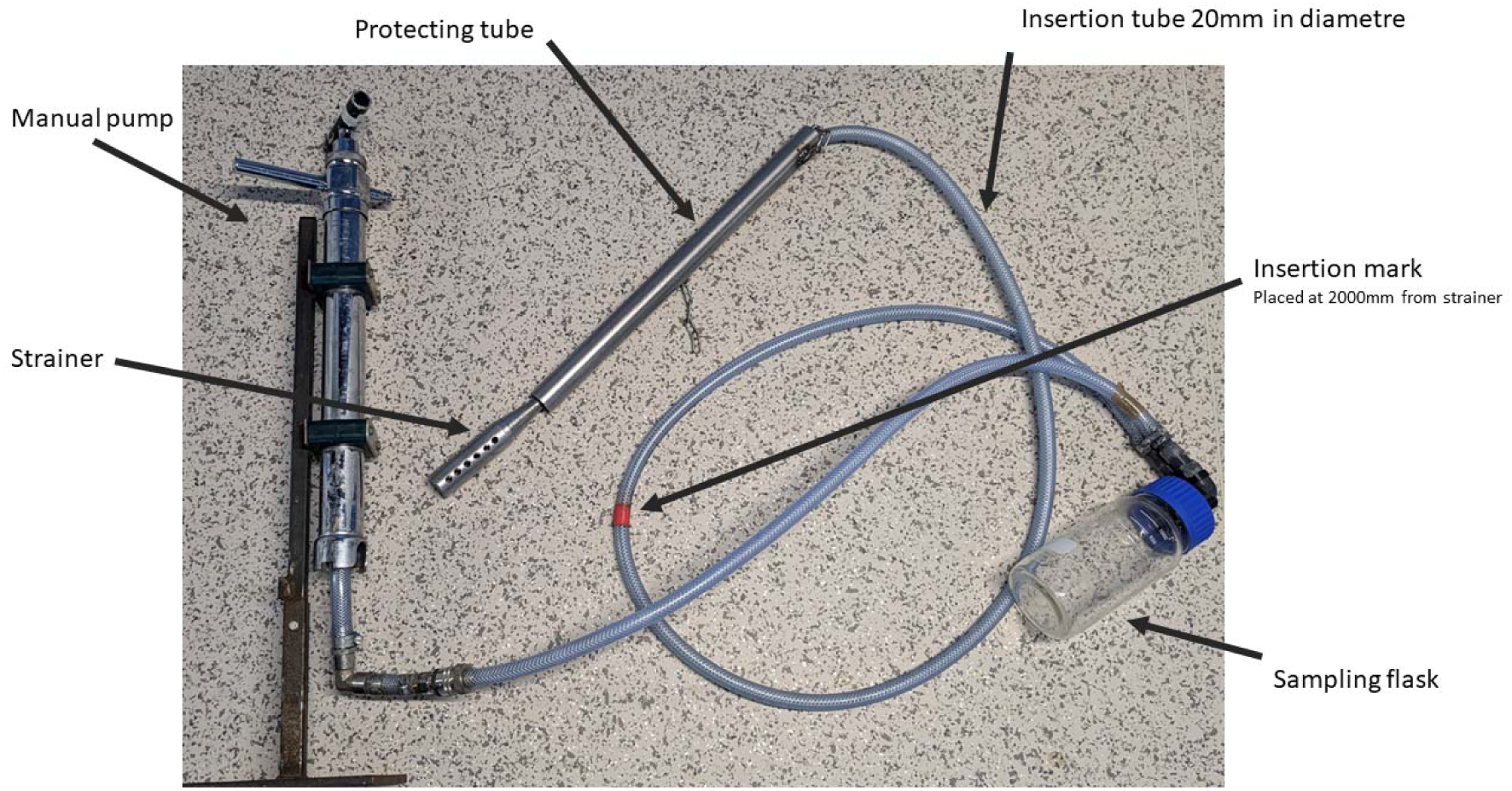
OSS sampling device used in the present study. The red mark on the tube was set at 200 cm from the strainer to ensure the placement in the ventral sac more than in the reticulum, as already described by Shen et al (2012).

**Fig. 2.**
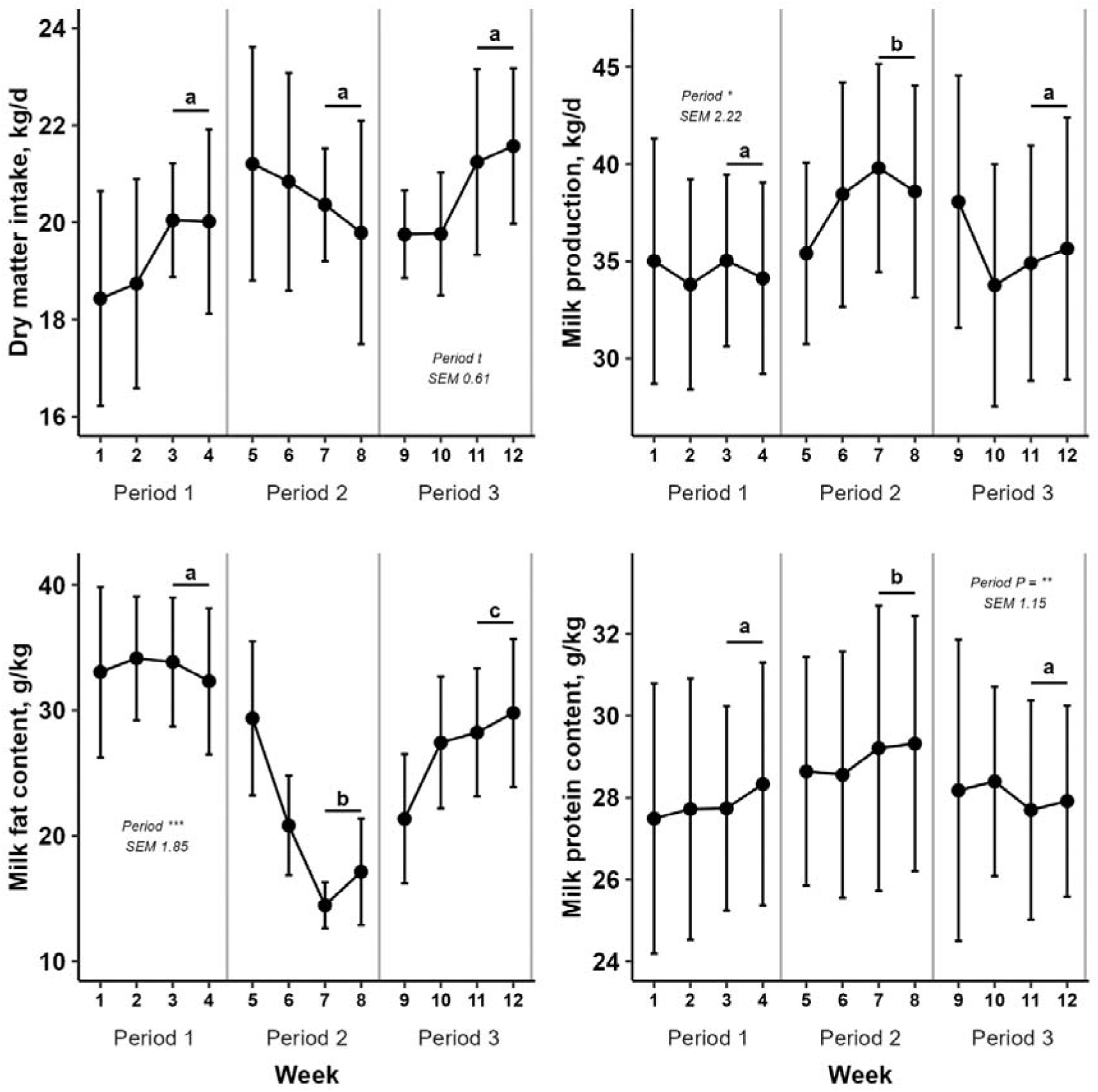
Dynamics of dry matter intake, milk production, milk fat and protein content throughout the experiment (mean ± SD, the statistical unit for statistical analyses was the average of the last two weeks for each period per cow, lsmeans are provided in table S1, period 1 includes weeks 1 to 4, period 2 weeks 5 to 8 and period 3 weeks 9 to 12). Lower case letters indicate significant differences between means by period. Statistical difference was achieved when P < 0.05

### Ruminal pH

Ruminal pH obtained through cannula sampling at 0830 h, 1330 h and 1630 h was on average for the 3 locations, 6.84, 5.82 and 6.32, respectively (Figure 3, Table S2). Whatever the sampling method or location, ruminal pH was lower in period 2 compared to periods 1 and 3, at both 1330 h and 1630 h (P <0.001). Between period 1 and 2, it dropped by 0.64 points at 1330 h (6.21 in period 1 vs. 5.57 in period 2) and by 0.17 points at 1630 h (6.34 in period 1 vs. 6.17 in period 2). At 0830 h, ruminal pH was lower in period 1 than in periods 2 and 3 (-0.22 points, P < 0.001), but in the 3 periods remained between 6.81 and 7.04. At both 0830 h and 1330 h, ruminal pH was higher when obtained by OSS than when sampled through the cannula, whatever the sampling method or location (P < 0.001). The pH difference was -0.44 points at 0830 h (7.28 for OSS and 6.84 on average for RCn, MCn and VSCn) and -0.56 points at 1630 h (6.38 for OSS and 5.82 on average for RCn, MCn and VSCn). No interaction between sampling location and method X period was observed.

**Fig. 3.**
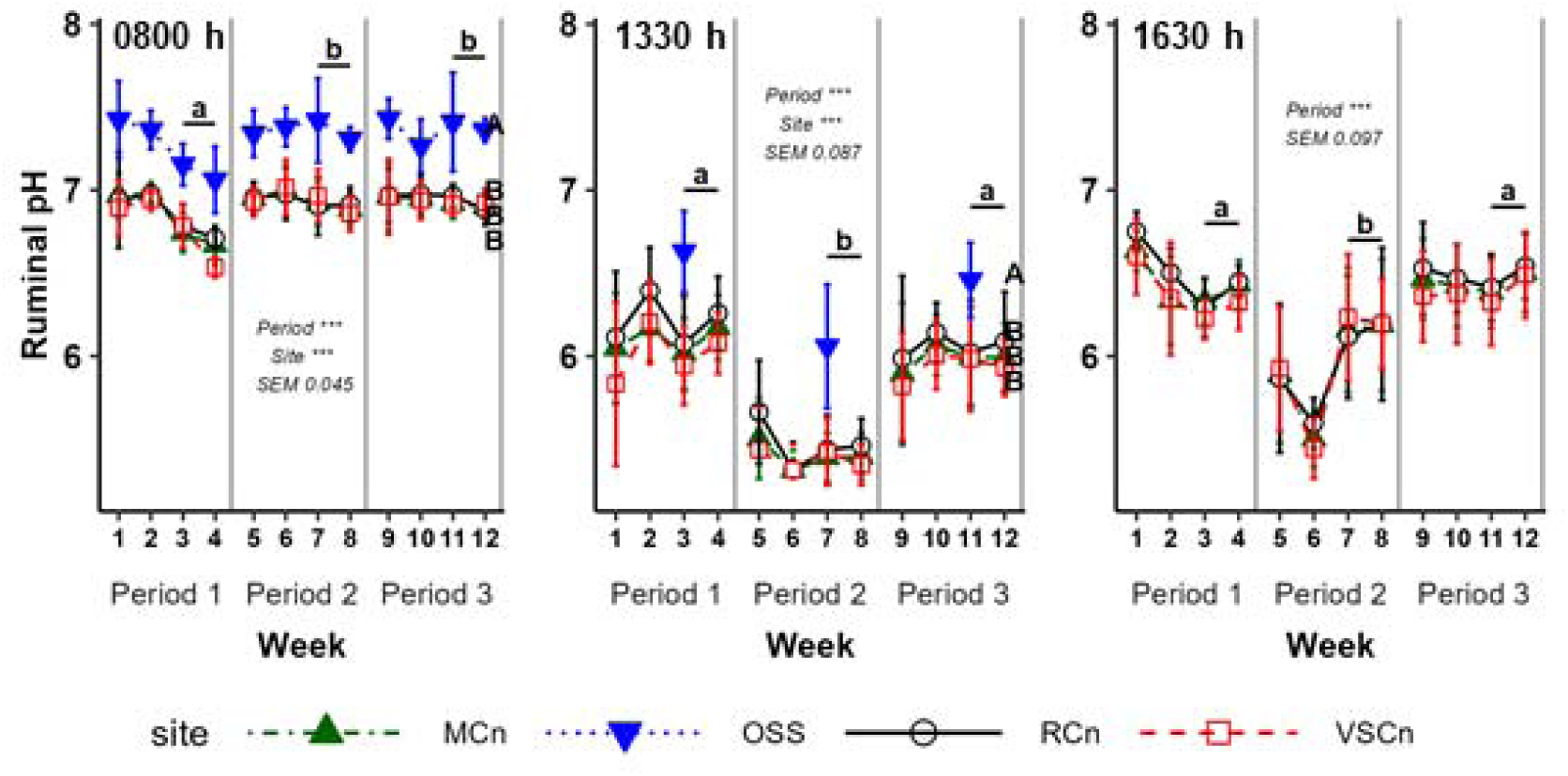
Dynamics of dry ruminal pH throughout the experiment (mean pH unit ± SD), the statistical unit for statistical analyses was the average per cow of the last two weeks for each period, lsmeans are provided in table S1, period 1 includes weeks 1 to 4, period 2 weeks 5 to 8 and period 3 weeks 9 to 12). Lower case letters indicate significant differences between means by period and upper-case letters indicate significant differences between means by sampling method/location. Statistical difference was achieved when P < 0.05

### Ruminal VFAs

Ruminal VFA concentration was on average for the 3 methods x 3 periods x 2 sampling times, 96.6 mmol/L at 0830 h and 172.9 mmol/L at 1330 h (Figures 4 and 5, Tables S3 and S4). At 0830 h, it was lower (P < 0.001) in period 2 (79.1 mmol/L) than in periods 1 and 3 (90.3 on average). At 1330 h, it increased (P < 0.001) between periods 1 (149 mmol/L) and 2 (188 mmol/L) and did not differ between periods 2 and 3 (184.7 mmol/L on average). At both 0830 h and 1330 h, ruminal VFA concentration was lower when obtained by OSS than when sampled through the cannula. At 0830 h, the differences between the methods of sampling was 27.0 mmol/L, with a value of 68.6 for OSS and 95.6 on average for VSCn and RCn. At 1330 h, the difference was 40.0 mmol/L, with a value of 146.2 for OSS and 166.2 on average for VSCn and RCn. At 0830 h, the ruminal VFA concentration was also slightly higher when sampled via the cannula in the ventral sac than in the reticulum (99.8 mmol/L for VSCn vs. 91.3 for RCn, P < 0.001) and the differences in ruminal VFA concentrations tended to be higher in period 1 than in periods 2 and 3 (interaction P < 0.10).

**Fig. 4.**
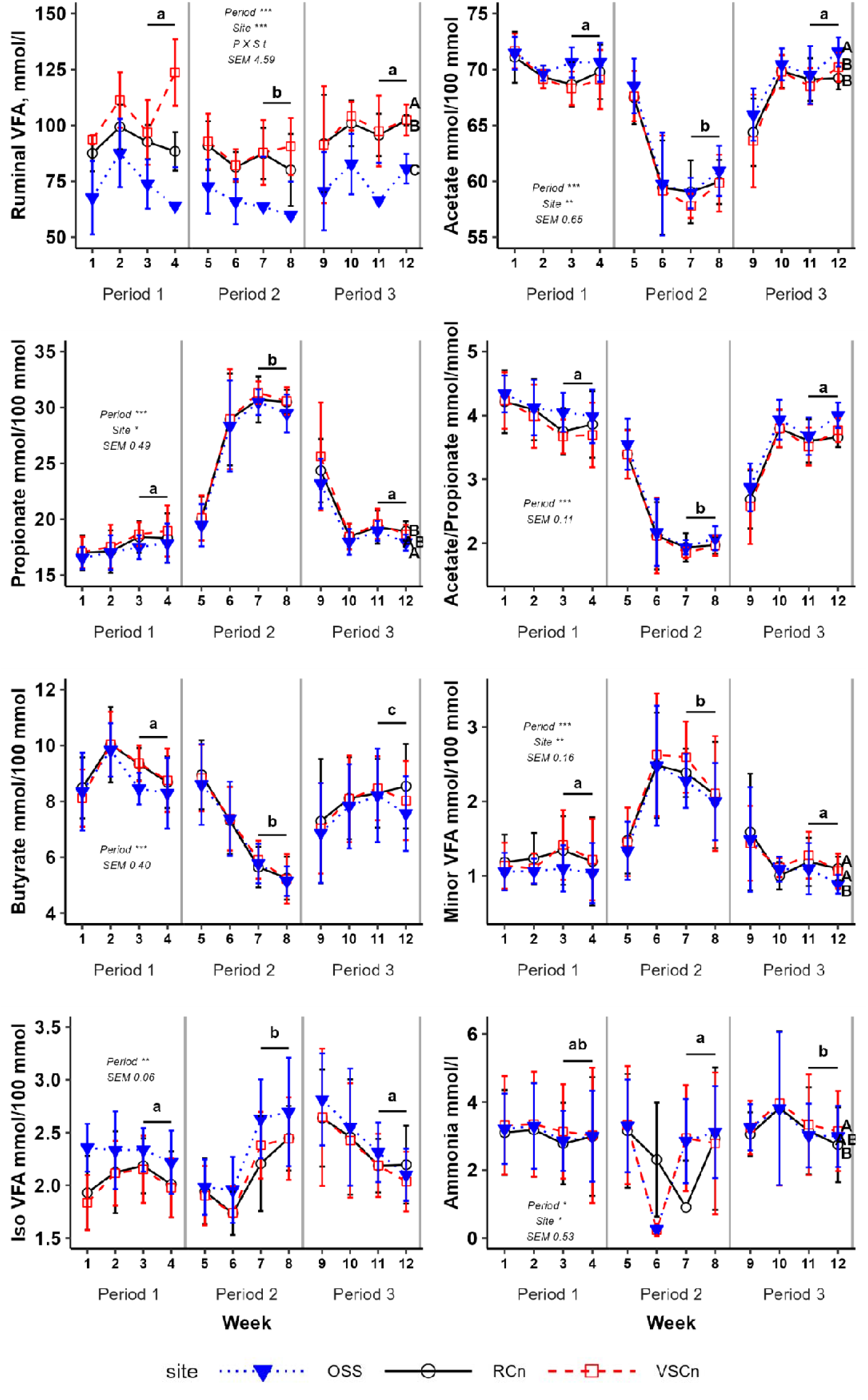
Dynamics of ruminal concentrations of volatile fatty acids (VFAs) and ammonia and VFA molar proportions at 0830 h throughout the experiment (mean ± SD), the statistical unit for statistical analyses was the average per cow of the last two weeks for each period, lsmeans are provided in table S3, period 1 includes weeks 1 to 4, period 2 weeks 5 to 8 and period 3 weeks 9 to 12). Lower case letters indicate significant differences between means by period and upper-case letters indicate significant differences between means by sampling method/location. Statistical difference was achieved when P < 0.05

**Fig. 5.**
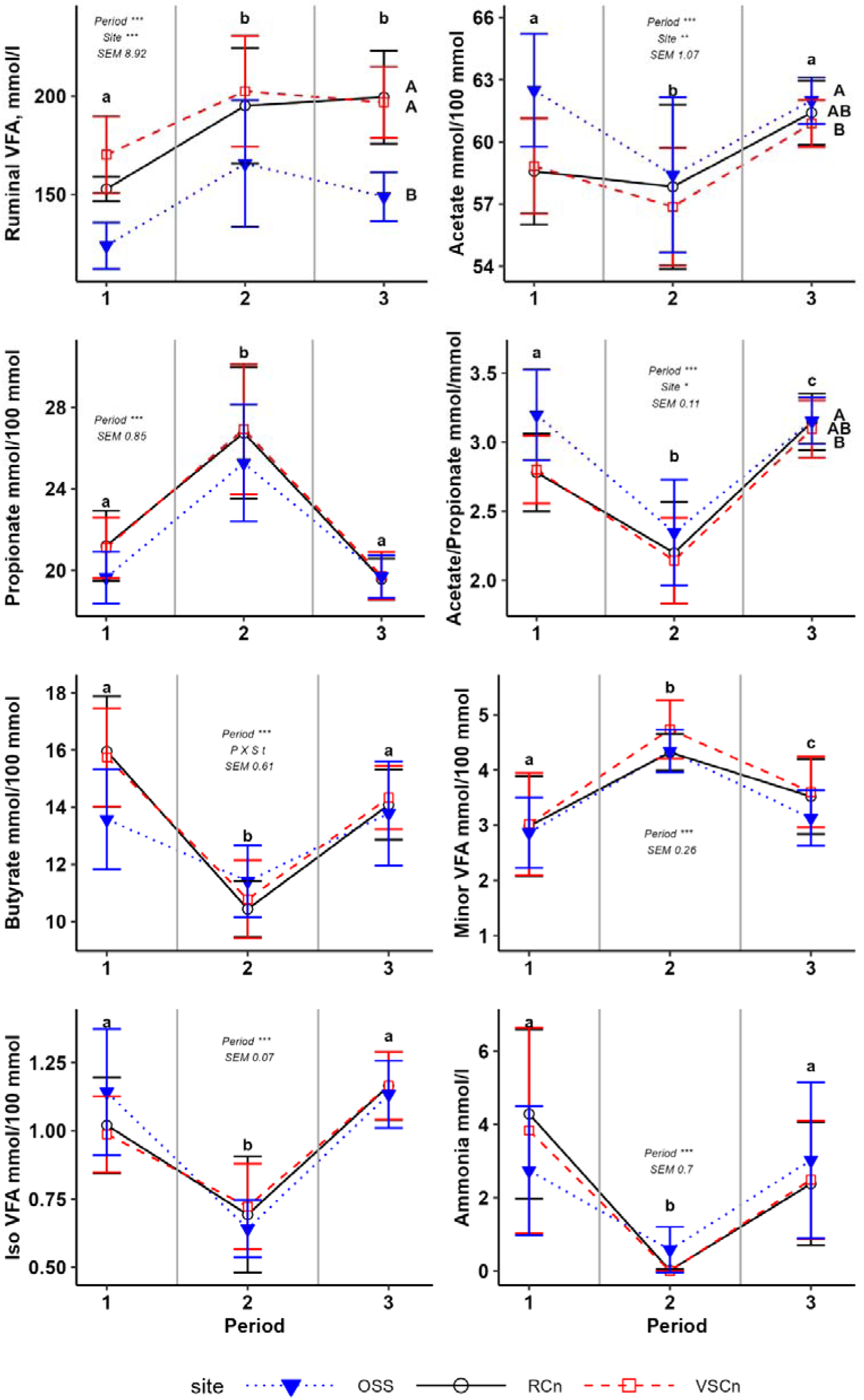
Dynamics of ruminal concentrations of volatile fatty acids (VFAs) and ammonia and VFA molar proportions at 1330 h throughout the experiment (mean ± SD), the statistical unit for statistical analyses was the average per cow of the last two weeks for each period, lsmeans are provided in table S3, period 1 includes weeks 1 to 4, period 2 weeks 5 to 8 and period 3 weeks 9 to 12). Lower case letters indicate significant differences between means by period and upper-case letters indicate significant differences between means by sampling method/location. Statistical difference was achieved when P < 0.05

Molar proportions of acetate, propionate, butyrate, minor and iso-VFAs were on average 66.1, 22.6, 7.5, 1.53 and 2.26 mmol/100 mmol at 0830 h, and 59.7, 22.2, 13.3, 3.61 and 0.96 mmol/100 mmol at 1330 h, respectively. Molar ratio acetate/propionate was on average 3.16 at 0830 h and 2.76 at 1330 h. At 0830 h, between periods 1 and 2, the molar proportions of acetate and butyrate, and the molar ratio acetate/propionate dropped considerably (- 10.3 and - 3.20 mmol/100 mmol, and - 1.89 mmol/mmol, respectively; P < 0.001), whereas the molar proportion of propionate and minor VFAs greatly increased (+ 3.84 and + 1.06 mmol/100 mmol; P < 0.001), with a similar trend for iso-VFAs (+ 0.30 mmol/100 mmol; P < 0.01). In period 3, molar proportions of acetate, propionate, minor VFAs and iso-VFAs, as well as the ratio between acetate and propionate, returned to values non-significantly different from those observed in period 1. Only the molar proportion of butyrate returned to a value slightly lower than that observed in period 1 (P<0.001).

At 1330 h, the same relative variation between periods 1 and 2 was observed, but with lower ranges of variation for the molar proportion of acetate and the molar ratio between acetate and propionate, and higher ranges of variation for molar proportions of propionate (+ 5.7 mmol/100 mmol), butyrate (- 4.23 mmol/100 mmol) and minor VFAs (+ 1.51 mmol/100 mmol). Only molar proportion of iso-VFAs decreased between periods 1 and 2 at 1330 h, whereas it increased at 0830 h. In period 3, molar proportions of acetate, propionate, butyrate and iso-VFAs also returned to values non-significantly different from those observed in period 1, whereas molar proportions of minor VFAs returned to a value slightly lower (P<0.001) than that observed in period 1 and the ratio between acetate and propionate to a value slightly higher at 1330 h (P<0.001).

Molar proportions of VFAs were differently affected by the sampling method or location according to the VFAs considered and the sampling time. At 0830 h, molar proportion of acetate was slightly higher in rumen fluid obtained by OSS than with the cannula on average for RCn and VSCn (+ 1.22 mol/100 mol, P < 0.001), whereas molar proportions of propionate and minor VFAs were slightly lower (- 0.8, P < 0.05 and - 0.20, P < 0.01, respectively). At 1330 h, molar proportion of acetate was higher in ruminal fluid obtained by OSS than in rumen fluid sampled through the cannula in VSCn (+ 2.1, P < 0.01). Similar variation was observed for acetate/propionate ratio (+ 0.22, P < 0.05). At 1330 h, molar proportion of butyrate tended to be lower in rumen fluid obtained by OSS than in rumen fluid sampled through the cannula in period 1 (interaction, P < 0.10).

### Ammonia

Ruminal concentration of ammonia was on average for the three methods 2.85 and 2.15 mmol/L at 0830 h and 1330 h, respectively. At 0830 h (Fig 4), it remained steady between periods 1 and 2 and increased between periods 2 and 3 (+ 0.55 mmol/L, P < 0.05), whereas at 1330 h (Fig 5) it greatly decreased between periods 1 and 2 (-3.42 mmol/L, P < 0.001) to return, in period 3, to a value close to that observed in period 1. At 0830 h, it was higher in VSCn than in RCn or in OSS samples (P < 0.05), whereas it was not affected (P > 0.10) by the sampling method or location at 1330 h. No interaction between sampling location and method X period was observed.

### Prediction of ventral sac fermentation parameters from OSS

Prediction of ruminal fermentation parameters in the ventral sac (pH, VFAs and NH3) from the OSS analysis are given in Figure 8. For all parameters (pH, ammonia and molar proportions of all VFAs), the determination coefficient ranged from r² = 0.70 to r² = 1. The RMSPE were low for the molar proportions of VFAs and relatively high for pH, ruminal concentrations of VFAs and ammonia. The RMSPE component due to the regression (ER) was very low for all parameters, ranging from 0% for acetate and ammonia molar proportion to 11% for molar proportion of iso-VFAs. All prediction slopes were either superposed with the bisector or presented a parallel offset for ruminal pH (+1.97) and ruminal VFAs (-0.32). For both pH and VFA concentration, the dominant component of the RMSPE was the ECT, which represented 80% of the RMSPE for pH and 69% for VFAs. For all molar proportions of VFAs the main component of RMSPE was the random error (EC) ranging from 57% for iso-VFAs to 84% for butyrate molar proportion.

### Ruminal concentrations of Ca, Mg, P, K and Na

At 0830 h, average ruminal concentrations for the 3 methods and the 3 periods of Ca, Mg, P, K and Na were 206.2, 42.8, 348.4, 797 and 2695 g/L, respectively (Figure 6 and Table S5). At 1330 h, these concentrations were 305.8, 104.8, 339.0, 1223 and 2265 g/L, respectively (Figure 7 and Table S6).

**Fig. 6.**
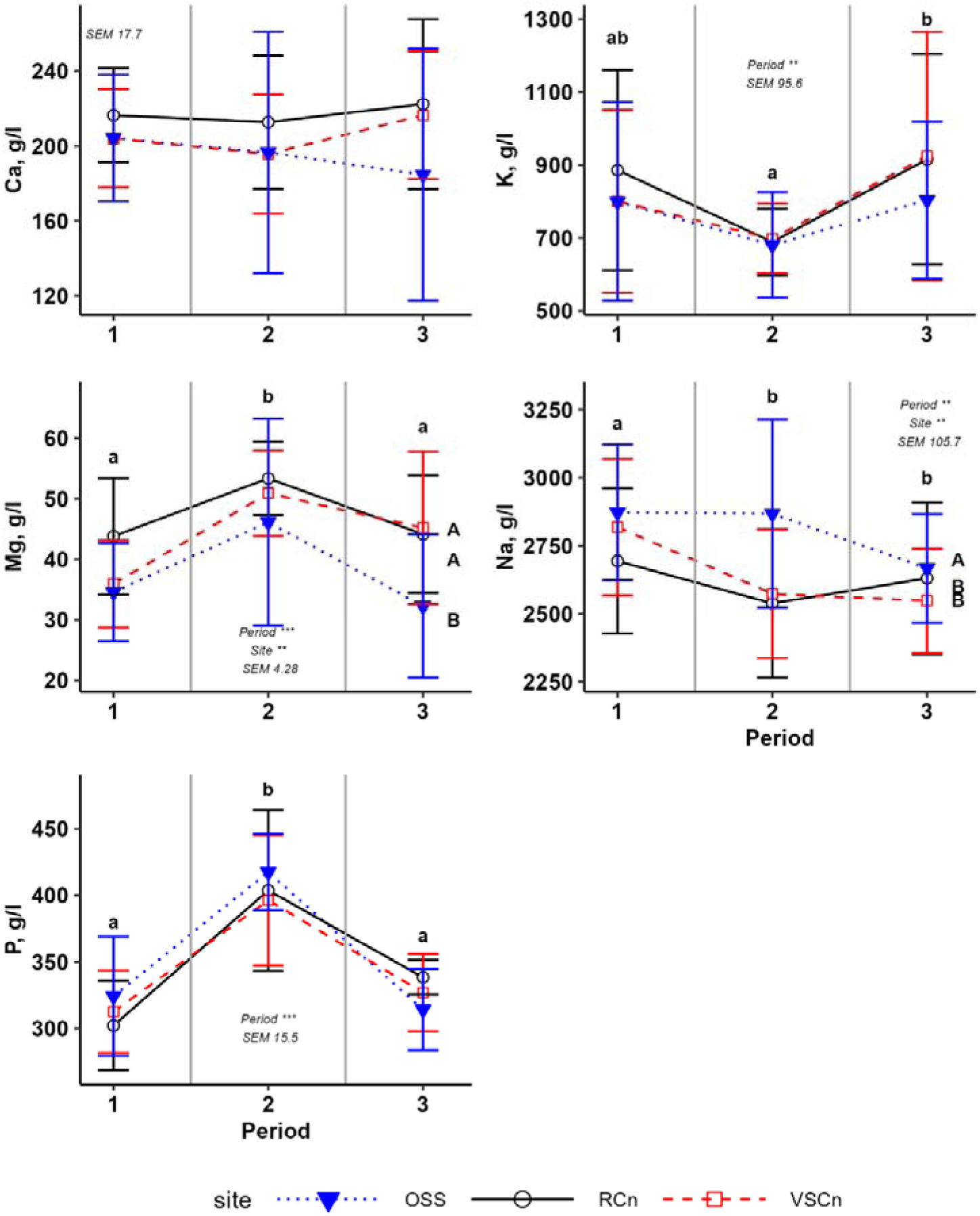
Dynamics of ruminal concentrations of minerals (Ca, Mg, P, K and Na) at 0830 h throughout the experiment (mean ± SD). Lower case letters indicate significant differences between means by period and upper-case letters indicate significant differences between means by sampling method/location. Statistical difference was achieved when P < 0.05

**Fig.7.**
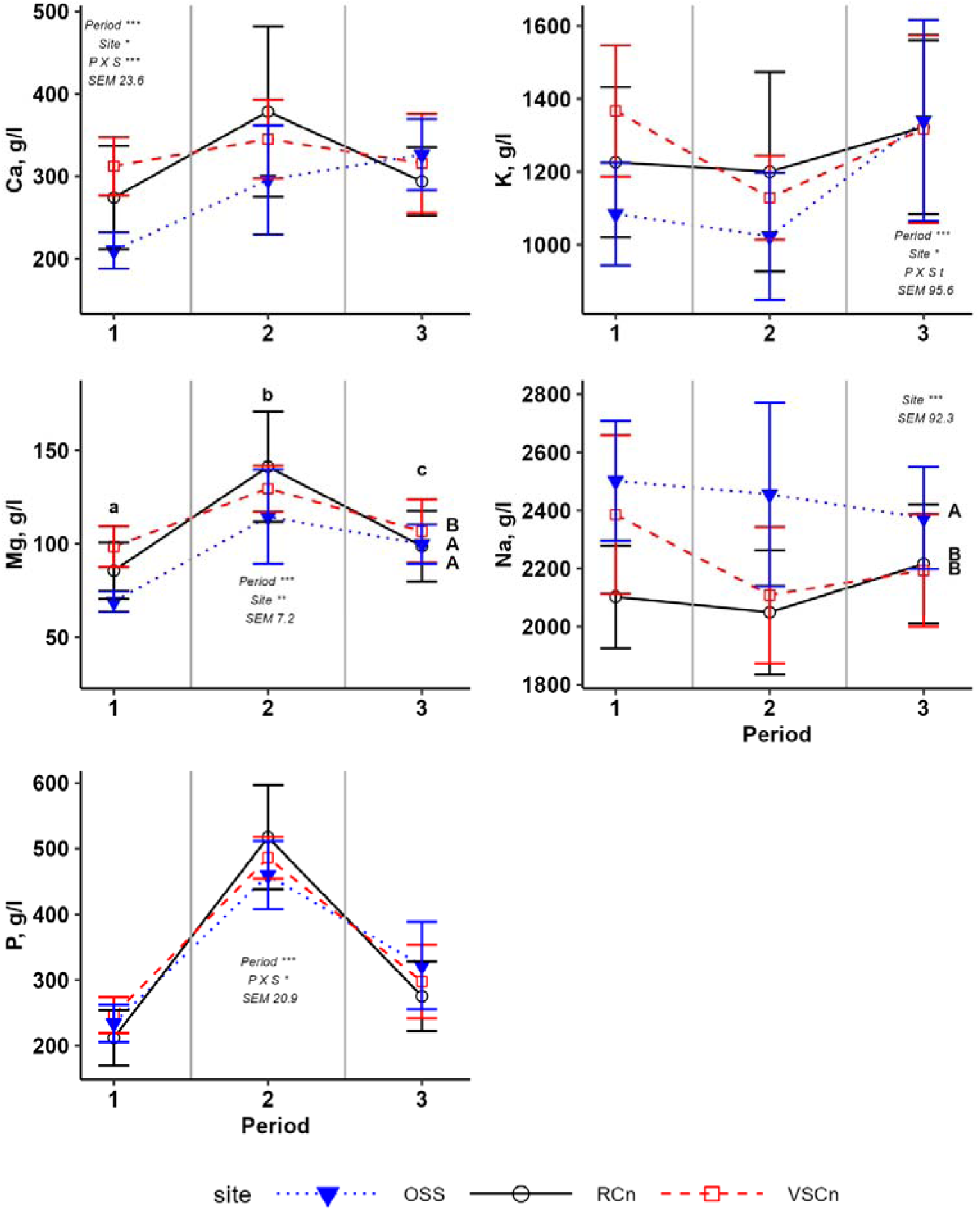
Dynamics of ruminal concentrations of minerals (Ca, Mg, P, K and Na) at 1330 h throughout the experiment (mean ± SD). Lower case letters indicate significant differences between means by period and upper-case letters indicate significant differences between means by sampling method/location. Letters are given when the interaction between period and method or sampling location was significant. Statistical difference was achieved when P < 0.05

**Fig.8.**
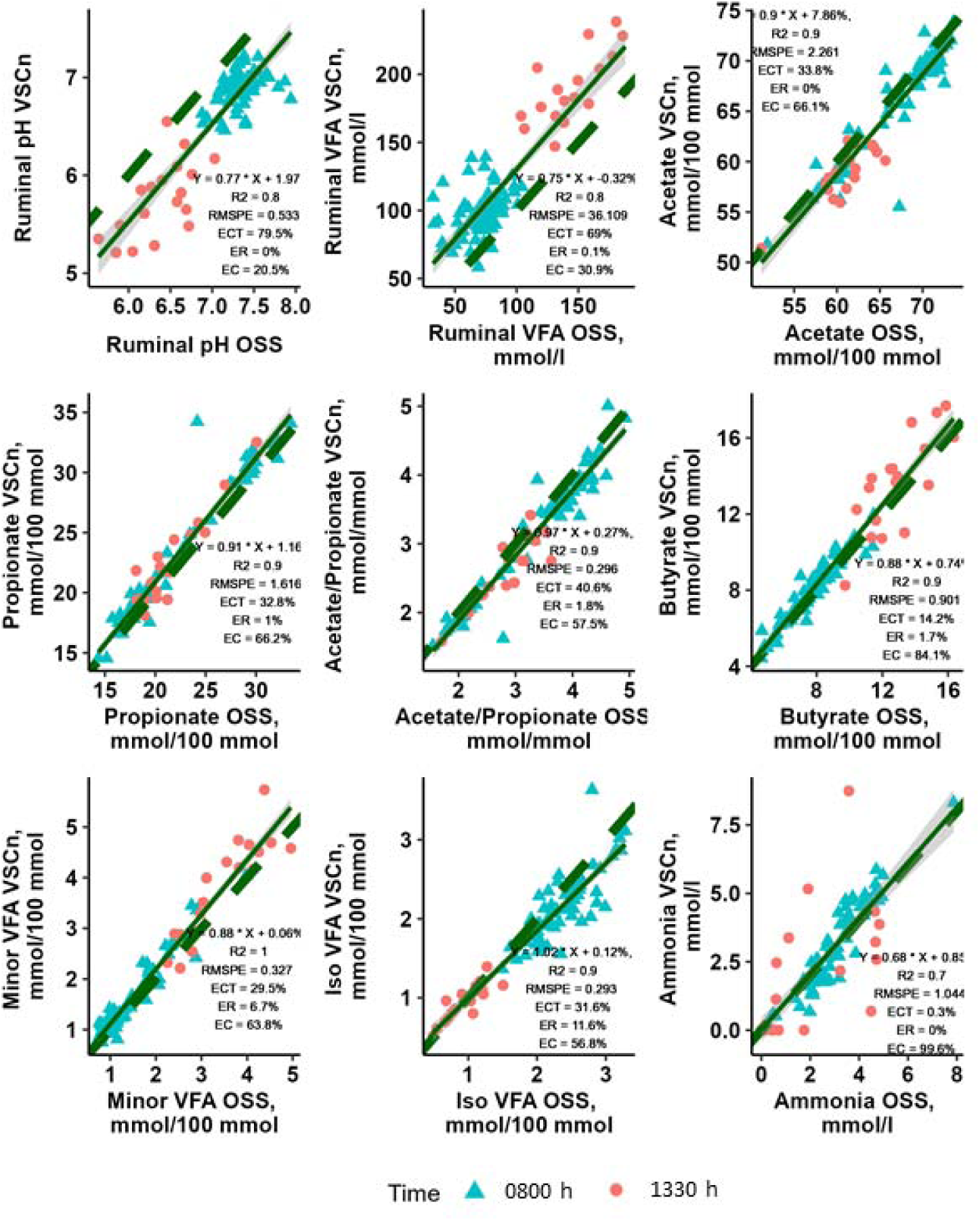
Linear regressions between sampling methods (OSS versus VSCn) for fermentation parameters (pH, total VFA, Major, minor and iso-VFA and ammonia). Linear regression adjustment coefficient is r², RMSPE gives the total error of prediction for each parameter whereas ECT, ER and EC, respectively, give error of central tendency, error due to regression and the random error component of RMSPE.

At 0830 h, ruminal concentrations of Mg and P increased between periods 1 and 2 and decreased in period 3 to a value close to that observed in period 1 (P < 0.001). Ruminal concentration of K was steady between periods 1 and 2 and increased between periods 2 and 3 (P < 0.01), whereas ruminal concentration of Na decreased between periods 1 and 2 and did not vary between periods 2 and 3 (P< 0.01).

Ruminal concentration of Mg was lower when ruminal fluid was obtained by OSS rather than sampled through the cannula (P < 0.01), whereas ruminal concentration of Na was higher (P < 0.01). Ruminal concentration of Ca was affected neither by the period nor by the sampling method or location (P > 0.10).

At 1330 h, as at 0830 h, ruminal concentration of Mg increased between periods 1 and 2 and decreased in period 3 to a value close to that observed in period 1 (P < 0.001). Ruminal concentration of Mg was also lower when ruminal fluid was obtained by OSS rather than sampled through the cannula (P < 0.01), whereas ruminal concentration of Na was also higher (P < 0.01). However, the ruminal concentrations of the other analyzed elements were all affected, or tended to be affected by the interaction between the period and the sampling method. Ruminal concentration of Ca was clearly lower when ruminal fluid was obtained by OSS rather than sampled through the cannula in periods 1 and 2, whereas it did not differ between the sampling methods in period 3 (P < 0.001). The same variations in tendency were observed for the ruminal concentration of K (P <0.10). As at 0830 h, ruminal concentration of P clearly increased between periods 1 and 2 and decreased in period 3 to a value close to that observed in period 1, but this increase was clearer when ruminal fluid was sampled through the cannula than obtained by OSS (P < 0.05).

## Discussion

In the present paper, our objective was to validate an optimized OSS method for its ability to highlight differences between treatments in a model experiment consisting of studying the effect of the inclusion of an acidogenic diet on the dynamics of ruminal composition in lactating dairy cows. The induction of the acidogenic diet was intended to increase variability in the analyses made on the various matrices sampled, such as the rumen fluid via OSS or through a fistula. Acidosis is a nutritional disease that does not have specific clinical signs, but many fermentation or production parameters are relevant for the onset of acidosis in ruminants (Calsamiglia et al., 2012). In the present study, the nutritional challenge based on the distribution of a high-energy diet successfully promoted acidosis in Period 2. All major VFAs were affected in Period 2 as expected: acetate was significantly decreased and propionate was significantly increased leading to a C2/C3 ratio that dropped below 3. Such a result has already been shown to be related to ruminal acidosis onset (Sauvant and Peyraud, 2010). These modifications in VFA profile induced the lowering of rumen pH. Rumen pH was altered, as expected, mostly after feeding at 1330 h, whereas no effect was observed at 0830 h. Such an observation is consistent with the description of Villot et al (2017) postulating that mean rumen pH does not necessarily decrease during bouts of acidosis, but is highly variable and increases in range during the day (Villot et al., 2017). Finally, the acidosis challenge led to a significant reduction of milk yield and milk quality, with a lowered fat/protein ratio. This result is consistent with what is often observed during acidosis challenges in experimental dairy cows (Silberberg et al., 2024). Altogether, those observations validated the nutritional acidotic challenge applied to the animals in the present experiments.

Our results showed higher ruminal pH and lower ruminal VFA concentrations with OSS compared to cannula sampling. This result is in line with many other studies that have performed the same comparison (Duffield et al., 2004; Shen et al., 2012; van Gastelen et al., 2019), although some studies did not observe any bias (Geishauser and Gitzel, 1996; Shen et al., 2012). The systematic pH bias we observed between OSS and cannula sampling was also within the range observed by several authors, + 0.48 in our case compared to + 0.34 for Duffield et al. (2004), + 0.32 for Shen et al. (2012) and + 0.55 for van Gastelen et al. (2019) under similar sampling conditions. However, the higher proportions of acetate and lower proportions of propionate and minor VFAs that we observed with OSS compared to VSCn sampling are specific to our study. Indeed, several studies have found that the sampling procedure did not affect the molar proportions of VFAs (Lodge-Ivey et al., 2009; van Gastelen et al., 2019; de Assis Lage et al., 2020). However, it should be noted that the differences we observed between OSS and VSCn were proportionally small: 2.04 % mol/100 mmol for C2, 4.18 % mol/100 mmol for C3 and 3.72 % mol/100 mmol for C4, compared to 7.14 percentage points for pH and 25.7% of mmol/L for VFAs. Although numerically small, we also observed a difference in VFA content between sampling sites in the rumen via the cannula, with slightly higher content in the ventral sac of the rumen than in the reticulum, which is consistent with literature data (Duffield et al., 2004; Shen et al., 2012). The purpose of comparing sampling sites was to account for differences in cannula sampling procedures between teams, although sampling in the ventral sac is recommended (Geishauser and Gitzel, 1996). Our results show that sampling at the bottom of the reticulum or in the ruminal ventral sac has little effect on rumen juice composition.

Our analyses of rumen mineral content confirmed that dilution by saliva is a probable reason for the differences in composition between rumen fluids obtained by OSS or cannula sampling. It has been found that the Na and P contents in saliva are higher than in rumen fluid, whereas the K content is lower (Bailey, 1961). Thus, in our experiment, the higher Na content of rumen fluid from OSS than in the cannula samples at 0830 h and 1330 h, and the lower K content at 1330 h, indicated a probable dilution by saliva of the samples obtained by OSS. This dilution also explains the lower Mg content of the rumen fluid from OSS than from cannula sampling and the same variations for Ca at 1330 h. This result is all the more consistent as the Ca and Mg levels observed in the rumen fluids from our experiment are high compared to those reported in ruminant saliva (McDougall, 1948; Riad et al., 1987; Dua and Care, 1998). Our results also confirm that saliva contamination is greater when samples are taken after a meal than before, as observed by others (Muizelaar et al., 2020). One question may be whether this dilution of samples obtained by OSS is directly due to the saliva included in the sample as the probe passes through the oesophagus or due to the sampling site being too dorsal for the probe, or both. The inclusion of saliva as the probe passes through the oesophagus can be reduced by eliminating the first 500 mL collected (Geishauser and Gitzel, 1996; Duffield et al., 2004; Shen et al., 2012), as we did in the present work. Dilution can also be caused by the sampling site being too close to the atrium, which is heavily irrigated with saliva during chewing. This can be controlled by checking that the probe is inserted deep enough to reach the rumen ventral sac (Shen et al., 2012). Although we took care to eliminate the first 500 mL sampled and checked that the probe reached the rumen ventral sac, we cannot draw firm conclusions about the exact causes of the salivary contamination we observed. The length of insertion of the tube in the oesophagus was checked to be 200 cm thanks to a mark made on the tube after preliminary tests on similar fistulated cows.

The question is whether sampling by OSS can be an alternative to rumen-fistulated animals in obtaining representative samples of ruminal fluid in an experimental protocol aimed at evaluating the effect of the inclusion of an acidogenic diet on the dynamics of ruminal composition in lactating dairy cows. Sampling from the ventral sac can be considered a benchmark for studying ruminal dynamics, as it is less subject to significant transient and immediate contamination by saliva associated with ingestion and rumination mastication (Shen et al., 2012). For all the parameters we measured, we observed a strong correlation and a slope closed to 1 between the measurements obtained by OSS and by cannula sampling in the rumen ventral sac (Geishauser and Gitzel, 1996), as well as a non-negligible but fixed bias for pH and VFA concentrations. This was consistent with the fact that the period effects were similar for all parameters regardless of the sampling method. This suggests that, although we are aware that the values obtained by OSS are not fully representative in absolute terms, sampling by OSS allows relative differences to be considered.

However, our regression curves also clearly showed very noisy predictions with OSS compared to cannula sampling for pH and VFA concentrations. For these two parameters, the predictive value of OSS sampling was even quite low if we look at the relationships within sampling times. Other authors have come to the same conclusions (Duffield et al., 2004). This shows that the predictive character of the analyses carried out on rumen content samples obtained by OSS is only guaranteed for large variations in pH, to be evaluated in relation to an average prediction error of OSS compared to cannula sampling of about 0.8 for pH and 36 mmol/L for VFA content. On the other hand, it is important to note that the prediction of VFA molar fractions remained generally satisfactory even within the sampling times. Finally, a solution to be favoured under experimental conditions could be to combine the use of telemetric ruminal boluses (Villot et al., 2018) for pH measurement with OSS sampling for the quantification of VFA molar proportions. Lastly, it is surprising that we did not observe any systematic bias related to a higher dilution of ammonia with OSS compared to cannula sampling, as for pH and ruminal VFA concentrations. We have no real explanation for this. However, given the limited variation in rumen protein balance (INRA, 2018) between our diets, the decrease in ammonia concentration with the acidogenic diets in period 2 was consistent with that of the iso-VFAs and with previously observed results comparing groups of animals at risk or not of acidosis (Golder et al., 2023).

Our results, as well as those of other publications, demonstrate the difficulty of obtaining a fully representative sample of ruminal contents by OSS, despite our efforts to follow the recommendations as closely as possible (Muizelaar et al., 2020). Our results also suggest that animal habituation, for example through medical training, may also be a parameter to consider. Indeed, we observed that after the meal, i.e. at the moment when sampling by OSS undoubtedly induces the greatest dilution by saliva, this dilution was probably reduced at the end of the experiment compared to the beginning, if we refer to the K and Ca contents of the rumen fluids. We also observed numerical variations in this direction for the Mg and Na contents of the rumen fluids. In general, the operators observed that OSS disturbed the animal more than cannula sampling. The effect of OSS on a certain number of behavioural indicators (Silberberg et al., 2015) and oesophageal lesions should be quantified. In our experiment, we limited the number of OSS to two per week. It would also be useful to know to what extent this number could be increased in animals undergoing medical training for this procedure. As we have seen before, another critical point is to determine the depth of insertion of the probe (Shen et al., 2012; Muizelaar et al., 2020). In the future, this operation may prove complicated in the absence of fistulated animals and it would undoubtedly be time to establish references between animal size and this parameter in order to refine the method.

## Conclusion

Our results show that OSS of rumen fluid is a relevant alternative to the use of fistulated cows for the quantification of VFA molar proportions. It may be an interesting alternative for measuring relative variations in ruminal pH and VFA concentrations, provided that the variability to be quantified is large enough, considering the random variations induced by dilution by saliva. Our results also illustrate that the procedure and recommendations for OSS still need to be refined, in particular by considering the use of trained animals or by specifying the recommendations for probe insertion depth. The relevance of OSS in quantifying the effect of acidogenic conditions on the microbiota and the relevance of OSS in terms of the quality of inoculum for gas test fermenters remain to be established.

## Ethics approval

All procedures were carried out in compliance with EU law on experimental animals and the ARRIVE guidelines. The experimental design and the experimental procedures were approved by a Regional Ethics Committee and the French Ministry of Research under number APAFiS #26894-2020081715322100_v2.

## Data and model availability statement

Data are available on request.

## Declaration of generative AI and AI-assisted technologies in the writing process

AI was only used to improve the English style (DeepL Write) before proofreading by a native English speaker.

## Acknowledgements

This research was supported by the INRAE Animal Physiology and Livestock Systems Division. The authors also thank ADISSEO, CARGILL, CMI-ROULLIER, LALLEMAND, MG2MIX, PHILEO-LESAFFRE, PROVIMI for funding and helpful discussion. They also thank Philippe Lamberton, Jean-Yves Thebault, Jean-Luc and Anthony Herouet for animal care and sampling on the experimental farm, Maryline Lemarchand, Séverine Urvoix, Maryvonne Texier at UMR PEGASE (INRAE) and Angélique Torrent at UMRH (INRAE) for sample treatment and analyses.

