## Supplemental tables for "Oral-stomach sampling to replace rumen-fistulated animals in ruminant nutrition research - a case study"

Table S1: Effect of the period on dry matter intake, milk production, milk fat and protein content of the six dairy cows involved in the experiment (the statistical unit was the average of the last two weeks for each period per cow, lsmeans are given)

|  | Period 1^1^ | Period 2 | Period 3 |  | SEM |  | P-value |
| --- | --- | --- | --- | --- | --- | --- | --- |
| Dry matter intake, kg/d | 20.0a | 20.1a | 21.4b |  | 0.60 |  | 0.055 |
| Milk production, kg/d | 34.6a | 39.2b | 35.3a |  | 2.20 |  | 0.016 |
| Milk fat content, g/kg | 33.1a | 15.8b | 29.0c |  | 1.90 |  | <0.001 |
| Milk protein content, g/kg | 28.0a | 29.3b | 27.8a |  | 1.20 |  | 0.009 |
| Ratio milk fat to protein contents, g/g | 1.18a | 0.54b | 1.04c |  | 0.05 |  | <0.001 |

^1^Periods 1 and 2: diet with low proportion of concentrate (12% DM) and low starch content (18% DM); period 2: diet with high proportion of concentrate (55% DM) and high starch content (29%DM). All periods lasted 4 weeks.

Table S2: Effect of the rumen location and methodology of rumen fluid sampling and period on ruminal pH of the six dairy cows involved in the experiment (the statistical unit was the average of the last two weeks for each period per cow, lsmeans are given)

|  | Loc.^1^ |  | Period 1^2^ | Period 2 | Period 3 |  | SEM |  | P_period | P_loc | P_inter |
| --- | --- | --- | --- | --- | --- | --- | --- | --- | --- | --- | --- |
| 08:30 | MCn |  | 6.71 | 6.9 | 6.91 |  | 0.045 |  | <0.001 | <0.001 | NS |
|  | RCn |  | 6.75 | 6.91 | 6.92 |  | 0.045 |  | <0.001 | <0.001 | NS |
|  | VSCn |  | 6.66 | 6.91 | 6.92 |  | 0.045 |  | <0.001 | <0.001 | NS |
|  | OSS |  | 7.11 | 7.35 | 7.39 |  | 0.045 |  | <0.001 | <0.001 | NS |
| 13:30 | MCn |  | 6.07 | 5.39 | 5.99 |  | 0.087 |  | <0.001 | <0.001 | NS |
|  | RCn |  | 6.14 | 5.44 | 6.04 |  | 0.087 |  | <0.001 | <0.001 | NS |
|  | VSCn |  | 5.99 | 5.39 | 5.97 |  | 0.087 |  | <0.001 | <0.001 | NS |
|  | OSS |  | 6.63 | 6.06 | 6.46 |  | 0.087 |  | <0.001 | <0.001 | NS |
| 16:30 | MCn |  | 6.38 | 6.16 | 6.44 |  | 0.093 |  | <0.001 | NS | NS |
|  | RCn |  | 6.37 | 6.14 | 6.48 |  | 0.093 |  | <0.001 | NS | NS |
|  | VSCn |  | 6.27 | 6.2 | 6.41 |  | 0.093 |  | <0.001 | NS | NS |

^1^Location and method of rumen sampling; RCn: reticulum through cannula; VSCn ventral sac through cannula; MCn: mix of reticulum liquor and ventral sac liquor (50/50, v/v) through cannula; OSS: oral-stomach sampling.

^2^Periods 1 and 2: diet with low proportion of concentrate (12% DM) and low starch content (18% DM); period 2: diet with high proportion of concentrate (55% DM) and high starch content (29%DM). All periods lasted 4 weeks.

Table S3: Effect of the rumen location and methodology of rumen fluid sampling and period on ruminal VFA concentrations, VFA composition and ammonia concentrations at 08:30 sampling for the six dairy cows involved in the experiment (the statistical unit was the average of the last two weeks for each period per cow, lsmeans are given)

|  | Loc^1^ |  | Period 1^2^ | Period 2 | Period 3 |  | SEM |  | P_period | P_loc | P_inter |
| --- | --- | --- | --- | --- | --- | --- | --- | --- | --- | --- | --- |
| Ruminal VFA^3^, mmol/L | RCn |  | 90.5 | 84.5 | 99.0 |  | 4.59 |  | <0.001 | <0.001 | 0.086 |
|  | VSCn |  | 110.2 | 89.3 | 99.9 |  |  |  |  |  |  |
|  | OSS |  | 68.9 | 63.4 | 73.5 |  |  |  |  |  |  |
| Acetate, mmol/100 mmol | RCn |  | 69.2 | 59.3 | 69.2 |  | 0.65 |  | <0.001 | 0.004 | NS |
|  | VSCn |  | 68.7 | 58.6 | 69.3 |  |  |  |  |  |  |
|  | OSS |  | 70.6 | 59.7 | 70.5 |  |  |  |  |  |  |
| Propionate, mmol/100 mmol | RCn |  | 18.4 | 30.6 | 19.1 |  | 0.49 |  | <0.001 | 0.023 | NS |
|  | VSCn |  | 18.8 | 31.0 | 19.1 |  |  |  |  |  |  |
|  | OSS |  | 17.7 | 30.0 | 18.4 |  |  |  |  |  |  |
| Acetate/Propionate, mmol/mmol | RCn |  | 3.81 | 1.95 | 3.63 |  | 0.107 |  | <0.001 | 0.009 | NS |
|  | VSCn |  | 3.68 | 1.90 | 3.64 |  |  |  |  |  |  |
|  | OSS |  | 4.02 | 2.00 | 3.84 |  |  |  |  |  |  |
| Butyrate, mmol/100 mmol | RCn |  | 9.0 | 5.6 | 8.4 |  | 0.40 |  | <0.001 | NS | NS |
|  | VSCn |  | 9.1 | 5.7 | 8.2 |  |  |  |  |  |  |
|  | OSS |  | 8.4 | 5.6 | 7.9 |  |  |  |  |  |  |
| Minor VFA, mmol/100 mmol | RCn |  | 1.27 | 2.29 | 1.14 |  | 0.164 |  | <0.001 | 0.007 | NS |
|  | VSCn |  | 1.32 | 2.40 | 1.17 |  |  |  |  |  |  |
|  | OSS |  | 1.07 | 2.15 | 0.99 |  |  |  |  |  |  |
| Iso VFA, mmol/100 mmol | RCn |  | 2.10 | 2.31 | 2.19 |  | 0.121 |  | 0.005 | NS | NS |
|  | VSCn |  | 2.06 | 2.40 | 2.11 |  |  |  |  |  |  |
|  | OSS |  | 2.28 | 2.64 | 2.21 |  |  |  |  |  |  |
| Ammonia, mmol/L | RCn |  | 2.88 | 1.70 | 2.95 |  | 0.530 |  | 0.041 | 0.035 | NS |
|  | VSCn |  | 3.07 | 2.90 | 3.24 |  |  |  |  |  |  |
|  | OSS |  | 2.92 | 2.95 | 3.00 |  |  |  |  |  |  |

^1^Location and method of rumen sampling; RCn: reticulum through cannula; VSCn ventral sac through cannula; MCn: mix of reticulum liquor and ventral sac liquor (50/50, v/v) through cannula; OSS: oral-stomach sampling.

^2^Periods 1 and 2: diet with low proportion of concentrate (12% DM) and low starch content (18% DM); period 2: diet with high proportion of concentrate (55% DM) and high starch content (29%DM). All periods lasted 4 weeks.

^3^VFA: volatile fatty acids

Table S4: Effect of the rumen location and methodology of rumen fluid sampling and period on ruminal VFA concentrations, VFA composition and ammonia concentrations at 13:30 sampling for the six dairy cows involved in the experiment (the statistical unit was the average of the last two weeks for each period per cow, lsmeans are given)

|  | Loc.^1^ |  | Period 1^2^ | Period 2 | Period 3 |  | SEM |  | P_period | P_loc | P_inter |
| --- | --- | --- | --- | --- | --- | --- | --- | --- | --- | --- | --- |
| Ruminal VFA^3^, mmol/L | RCn |  | 152.7 | 195.2 | 199.5 |  | 8.92 |  | <0.001 | <0.001 | NS |
|  | VSCn |  | 170.3 | 202.6 | 196.9 |  |  |  |  |  |  |
|  | OSS |  | 124.0 | 165.7 | 148.9 |  |  |  |  |  |  |
| Acetate, mmol/100 mmol | RCn |  | 58.6 | 57.8 | 61.4 |  | 1.07 |  | <0.001 | 0.037 | NS |
|  | VSCn |  | 58.8 | 56.9 | 60.9 |  |  |  |  |  |  |
|  | OSS |  | 62.5 | 58.4 | 62.0 |  |  |  |  |  |  |
| Propionate, mmol/100 mmol | RCn |  | 21.2 | 26.7 | 19.6 |  | 0.85 |  | <0.001 | NS | NS |
|  | VSCn |  | 21.1 | 26.9 | 19.7 |  |  |  |  |  |  |
|  | OSS |  | 19.6 | 25.3 | 19.7 |  |  |  |  |  |  |
| Acetate/Propionate, mmol/mmol | RCn |  | 2.78 | 2.20 | 3.15 |  | 0.117 |  | <0.001 | 0.026 | NS |
|  | VSCn |  | 2.80 | 2.14 | 3.10 |  |  |  |  |  |  |
|  | OSS |  | 3.20 | 2.35 | 3.15 |  |  |  |  |  |  |
| Butyrate, mmol/100 mmol | RCn |  | 16.0 | 10.4 | 14.1 |  | 0.61 |  | <0.001 | NS | 0.084 |
|  | VSCn |  | 15.7 | 10.8 | 14.3 |  |  |  |  |  |  |
|  | OSS |  | 13.6 | 11.4 | 13.8 |  |  |  |  |  |  |
| Minor VFA, mmol/100 mmol | RCn |  | 2.98 | 4.32 | 3.52 |  | 0.263 |  | <0.001 | NS | NS |
|  | VSCn |  | 3.02 | 4.74 | 3.60 |  |  |  |  |  |  |
|  | OSS |  | 2.87 | 4.34 | 3.13 |  |  |  |  |  |  |
| Iso VFA, mmol/100 mmol | RCn |  | 1.02 | 0.69 | 1.17 |  | 0.065 |  | <0.001 | NS | NS |
|  | VSCn |  | 0.99 | 0.72 | 1.17 |  |  |  |  |  |  |
|  | OSS |  | 1.14 | 0.64 | 1.13 |  |  |  |  |  |  |
| Ammonia, mmol/L | RCn |  | 4.29 | 0.02 | 2.38 |  | 0.702 |  | <0.001 | NS | NS |
|  | VSCn |  | 3.83 | 0.00 | 2.49 |  |  |  |  |  |  |
|  | OSS |  | 2.74 | 0.58 | 3.02 |  |  |  |  |  |  |

^1^Location and method of rumen sampling; RCn: reticulum through cannula; VSCn ventral sac through cannula; MCn: mix of reticulum liquor and ventral sac liquor (50/50, v/v) through cannula; OSS: oral-stomach sampling.

^2^Periods 1 and 2: diet with low proportion of concentrate (12% DM) and low starch content (18% DM); period 2: diet with high proportion of concentrate (55% DM) and high starch content (29%DM). All periods lasted 4 weeks.

^3^VFA: volatile fatty acids

Table S5: Effect of the rumen location and methodology of rumen fluid sampling and period on ruminal Na, K, Ca, P and Mg concentrations at 08:30 sampling for the six dairy cows involved in the experiment (the statistical unit was the average of the last two weeks for each period per cow, lsmeans are given)

|  | Loc^1^ |  | Period 1^2^ | Period 2 | Period 3 |  | SEM |  | P_period | P_loc | P_inter |
| --- | --- | --- | --- | --- | --- | --- | --- | --- | --- | --- | --- |
| Ruminal Na, g/L | RCn |  | 2694 | 2539 | 2631 |  | 105.7 |  | 0.007 | 0.009 | NS |
|  | VSCn |  | 2858 | 2574 | 2548 |  |  |  |  |  |  |
|  | OSS |  | 2873 | 2868 | 2667 |  |  |  |  |  |  |
| Ruminal K, g/L | RCn |  | 886 | 689 | 916 |  | 95.6 |  | 0.007 | NS | NS |
|  | VSCn |  | 775 | 698 | 924 |  |  |  |  |  |  |
|  | OSS |  | 800 | 681 | 804 |  |  |  |  |  |  |
| Ruminal Ca, g/L | RCn |  | 216.5 | 212.8 | 222.5 |  | 17.69 |  | NS | NS | NS |
|  | VSCn |  | 206.2 | 195.7 | 216.5 |  |  |  |  |  |  |
|  | OSS |  | 204.3 | 196.6 | 184.7 |  |  |  |  |  |  |
| Ruminal P, g/L | RCn |  | 302.1 | 403.7 | 338.3 |  | 15.52 |  | <0.001 | NS | NS |
|  | VSCn |  | 312.9 | 396.2 | 326.9 |  |  |  |  |  |  |
|  | OSS |  | 324.1 | 417.5 | 314.1 |  |  |  |  |  |  |
| Ruminal Mg, g/L | RCn |  | 43.8 | 53.4 | 44.2 |  | 4.28 |  | <0.001 | 0.005 | NS |
|  | VSCn |  | 34.9 | 50.9 | 45.2 |  |  |  |  |  |  |
|  | OSS |  | 34.6 | 46.2 | 32.3 |  |  |  |  |  |  |

^1^Location and method of rumen sampling; RCn: reticulum through cannula; VSCn ventral sac through cannula; MCn: mix of reticulum liquor and ventral sac liquor (50/50, v/v) through cannula; OSS: oral-stomach sampling.

^2^Periods 1 and 2: diet with low proportion of concentrate (12% DM) and low starch content (18% DM); period 2: diet with high proportion of concentrate (55% DM) and high starch content (29%DM). All periods lasted 4 weeks.

Table S6: Effect of the rumen location and methodology of rumen fluid sampling and period on ruminal Na, K, Ca, P and Mg concentrations at 13:30 sampling for the six dairy cows involved in the experiment (the statistical unit was the average of the last two weeks for each period per cow, lsmeans are given)

|  | Loc^1^ |  | Period 1^2^ | Period 2 | Period 3 |  | SEM |  | P_period | P_loc | P_inter |
| --- | --- | --- | --- | --- | --- | --- | --- | --- | --- | --- | --- |
| Ruminal Na, g/L | RCn |  | 2103 | 2049 | 2216 |  | 92.3 |  | NS | <0.001 | NS |
|  | VSCn |  | 2386 | 2108 | 2193 |  |  |  |  |  |  |
|  | OSS |  | 2502 | 2456 | 2374 |  |  |  |  |  |  |
| Ruminal K, g/L | RCn |  | 1226 | 1200 | 1323 |  | 87.4 |  | <0.001 | 0.031 | 0.074 |
|  | VSCn |  | 1367 | 1129 | 1318 |  |  |  |  |  |  |
|  | OSS |  | 1084 | 1023 | 1341 |  |  |  |  |  |  |
| Ruminal Ca, g/L | RCn |  | 274.3 | 378.6 | 293.9 |  | 23.66 |  | <0.001 | 0.01 | 0.009 |
|  | VSCn |  | 312.3 | 345.2 | 315.8 |  |  |  |  |  |  |
|  | OSS |  | 210.0 | 295.9 | 326.4 |  |  |  |  |  |  |
| Ruminal P, g/L | RCn |  | 211.8 | 517.5 | 275.3 |  | 20.93 |  | <0.001 | NS | 0.044 |
|  | VSCn |  | 246.7 | 486.2 | 297.6 |  |  |  |  |  |  |
|  | OSS |  | 234.1 | 459.6 | 322.0 |  |  |  |  |  |  |
| Ruminal Mg, g/L | RCn |  | 85.6 | 141.2 | 98.7 |  | 7.19 |  | <0.001 | 0.007 | NS |
|  | VSCn |  | 98.4 | 129.5 | 106.8 |  |  |  |  |  |  |
|  | OSS |  | 69.0 | 114.5 | 99.8 |  |  |  |  |  |  |

^1^Location and method of rumen sampling; RCn: reticulum through cannula; VSCn ventral sac through cannula; MCn: mix of reticulum liquor and ventral sac liquor (50/50, v/v) through cannula; OSS: oral-stomach sampling.

^2^Periods 1 and 2: diet with low proportion of concentrate (12% DM) and low starch content (18% DM); period 2: diet with high proportion of concentrate (55% DM) and high starch content (29%DM). All periods lasted 4 weeks.
